# SNP calling, haplotype phasing and allele-specific analysis with long RNA-seq reads

**DOI:** 10.1101/2025.05.26.656191

**Authors:** Neng Huang, Human Pangenome Reference Consortium, Heng Li

## Abstract

Long-read RNA sequencing (lrRNA-seq) is a powerful technology to link transcript structures to genetic variants but such analysis is not often performed due to the lack of end-user tools. Here, we introduce longcallR for joint SNP calling, haplotype phasing, and allele-specific analysis, which achieves high accuracy on benchmark datasets. Applied to 202 human samples, longcallR identified 88 significant allele-specific splicing events per sample on average. 46% of them involved unannotated junctions.

---

lrRNA-seq technologies, including Iso-Seq and MAS-Seq from Pacific Biosciences (PacBio) and cDNA sequencing and direct-RNA sequencing (dRNA) from Oxford Nanopore Technologies (ONT), are gaining interest in transcriptomic research for the potential to capture full-length transcripts. This capability provides a comprehensive view of gene expression, transcript quantification^1–7^ and alternative splicing^8,9^. As individual lrRNA-seq reads can span multiple single nucleotide polymorphisms (SNP), they are more effective for haplotype phasing and enable the study of allele-specific events which can help us understand the complex relationships among genetic variation, splicing disruptions, and disease.

Allele-specific analyses require accurate SNP calling. While calling SNPs from long genomic reads has been well established^10–14^, calling from lrRNA-seq data is faced with unique challenges, including uneven read coverage due to varying gene expression, frequent alignment errors around splice sites, skewed allele fraction caused by allele-specific expression, and post-transcriptional RNA editing events that may be mistaken as SNPs. Among the few SNP calling pipelines developed for lrRNA-seq^15,16^, only Clair3-RNA^17^ works with Nanopore data but it does not phase SNPs or perform allele-specific analyses.

Here, we present longcallR for SNP calling, haplotype phasing, and allele-specific analysis using lrRNA-seq data. LongcallR consists of three modules. First, longcallR-nn utilizes a deep convolutional neural network^18^ to analyze pileup images generated from read-to-reference alignments, predicting the genotype and zygosity of SNPs (**Fig. 1a**, Methods). The basic principle is similar to Clair3-RNA, but the model architecture, the feature sets and the training data construction all differ. Second, implemented in the Rust programming language, longcallR-phase uses a probabilistic model to jointly refine SNP calls and perform haplotype phasing (**Fig. 1b**, Methods). It also functions as a standalone algorithm, directly producing phased SNP calls from lrRNA-seq alignment. The third module of longcallR tests allele-specific expression (ASE) and allele-specific junctions (ASJ) from a single sample or jointly across multiple samples (**Fig. 1c**, Methods).

**Fig. 1.**
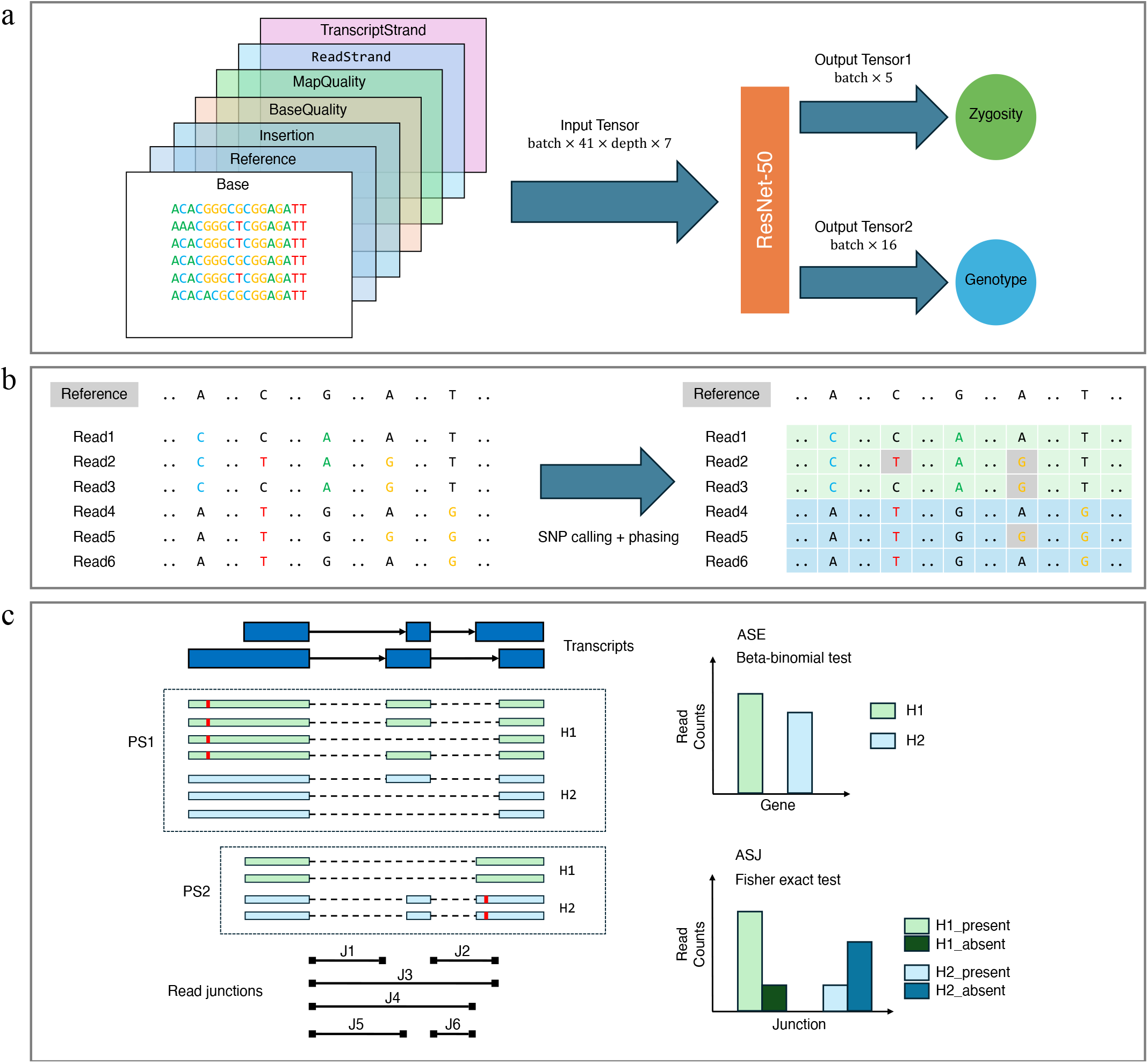
Overview of the longcallR algorithm. **A**, the longcallR-nn module for RNA SNP calling. It constructs a 7-channel image representing a 41 bp flanking region around each candidate SNP (left). This image is processed by a ResNet-50 convolutional neural network, which outputs two classifications (right): zygosity (0/0, 0/1, 1/1, ½, and A-to-I RNA editing) and genotype (combinations of reference alleles {A, C, G, T} including alternate alleles {A, C, G, T}). **B**, the longcallR-phase module for SNP filtering and haplotype phasing. It extracts read alleles at candidate heterozygous SNP sites (left) and then iteratively refines read haplotags (H1 or H2) and SNP haplotypes to minimize conflicts between read alleles and haplotypes. The phased reads (green: H1, blue: H2) are then used to distinguish true SNPs, sequencing errors, and A-to-I RNA editing events (right). **C**, allele-specific analysis using longcallR. Phased reads are assigned to the closest gene based on the longest exonic matching length, and read junctions are extracted from sequencing data (left). For allele-specific expression analysis (top-right), the module quantifies haplotype-specific read counts and applies a beta-binomial test. For allele-specific junction analysis (bottom-right), the module separately counts junction inclusion and exclusion reads for each haplotype and uses Fisher’s exact test to assess haplotype-specific splicing differences. The Benjamin-Hochberg procedure is applied for controlling the false positive rate.

We evaluated the SNP calling accuracy of longcallR and Clair3-RNA on 12 human datasets sequenced using PacBio Mas-Seq, Iso-Seq, as well as Nanopore cDNA and dRNA (**Supplementary Table 1**). We take genomic SNPs from Genome-In-A-Bottle (GIAB) or from short-read SNP calls as the ground truth (Methods). For the PacBio datasets (**Fig. 2a, Supplementary Table 2**), longcallR achieved precision ranging from 98.5% to 99.0% and recall from 91.0% to 95.3% across HG002, HG004, and HG005, the GIAB samples. It consistently outperformed Clair3-RNA in precision by 1.4% on average, though Clair3-RNA excelled in sensitivity. This is partly because longcallR requires SNPs in phase, which helps to reduce false positives but may miss true SNPs due to phasing difficulties in rare cases. The F1 scores of longcallR and Clair3-RNA were comparable at 95.7%. Both callers achieved higher sensitivity on the Mas-Seq datasets than on the Iso-Seq datasets, owning to the higher Mas-Seq coverage. Nanopore SNP calling is more challenging due to the higher sequencing error rate (**Fig. 2b, Supplementary Table 2**). The precision and recall of longcallR dropped to 85.8–98.0% and 73.9–87.3%, respectively, across GIAB samples. LongcallR again achieved higher precision at the cost of lower sensitivity in comparison to Clair3-RNA. Among different Nanopore data types, longcallR had the highest accuracy on two recent dRNA (RNA004) datasets. The SNP calling accuracy is similar in exonic regions (**Supplementary Table 3**). Clair3-RNA can call insertions and deletions in addition to SNPs. It reached a precision of 82.7% for HG004:MAS:cDNA and 60.5% for HG004:ONT:cDNA, much lower than the precision of SNP calling. This high error rate will complicate downstream allele-specific analysis, so we only focused on SNPs with longcallR.

**Fig. 2.**
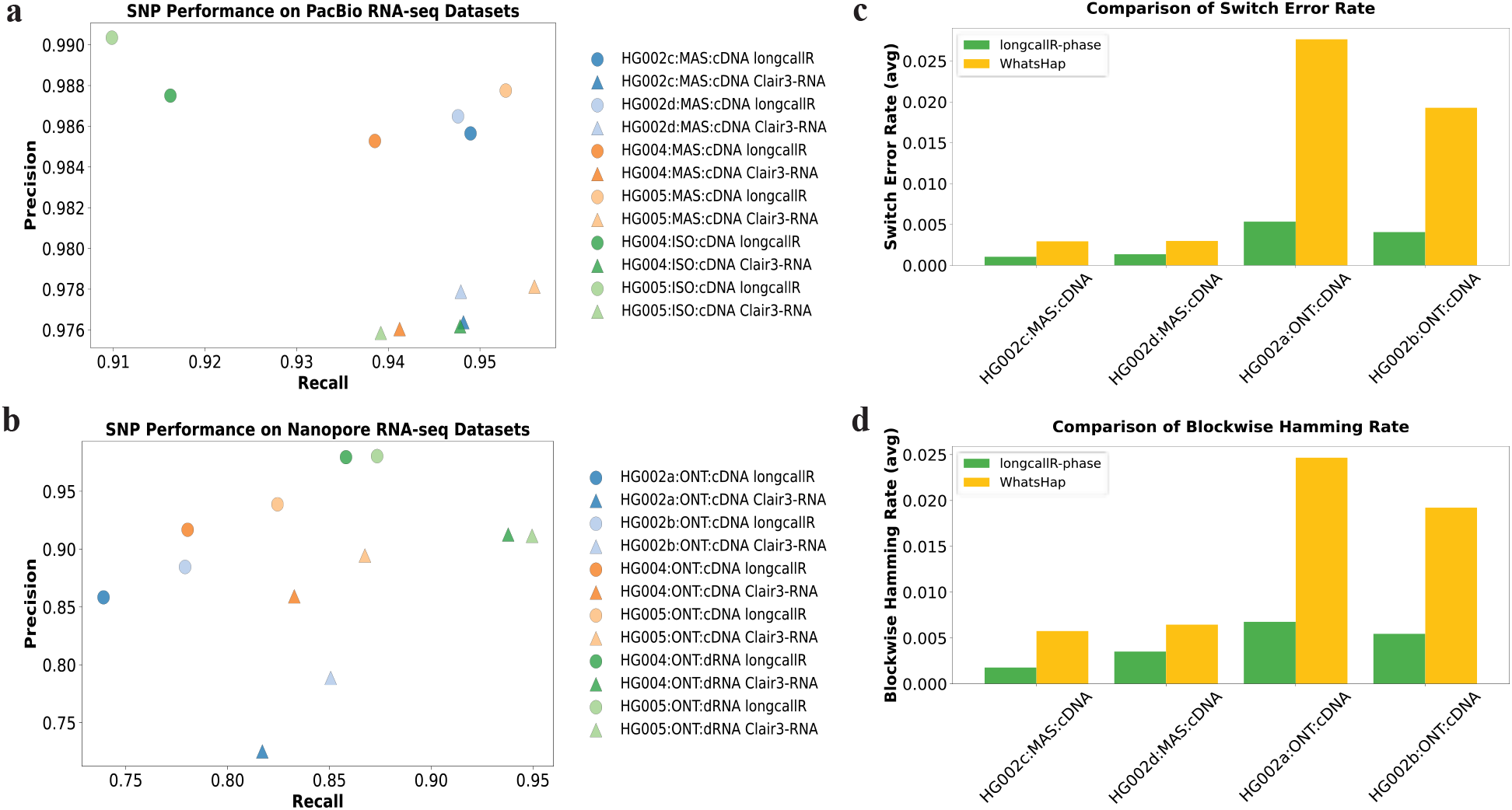
Accuracy of SNP calling and haplotype phasing. **A**, precision and recall for SNP calling on PacBio datasets between longcallR (circles) and Clair3-RNA (triangles). Colors represent different samples. HG002 genomic SNPs from GIAB are taken as the ground truth. **B**, precision and recall for SNP calling on Nanopore datasets. **C**, Phasing switch error rates between longcallR-phase (green) and WhatsHap (yellow) across PacBio and Nanopore datasets. Trio-based HG002 SNP phasing from GIAB is taken as the ground truth. Lower switch error rates indicate higher phasing accuracy. **D**, Phasing hamming error rates in haplotype phasing.

LongcallR also works as a standalone phasing tool for lrRNA-seq data. To evaluate its phasing accuracy, we compared longcallR to WhatsHap on the HG002c:MAS:cDNA and HG002b:ONT:cDNA datasets, taking phased SNPs from GIAB^19^ as the ground truth. Given SNPs called by longcallR, longcallR produced longer phased blocks at lower phasing switch and hamming error rates (**Fig. 2c and 2d, Supplementary Table 5**). To reduce potential biases in calling algorithms, we also provided Clair3-RNA SNPs as input. LongcallR still produced lower phasing error rates with comparable or longer phase block lengths (**Supplementary Table 6**). Notably, both longcallR and WhatsHap generated longer phased blocks on longcallR SNP calls than on Clair3-RNA calls, especially on the Nanopore datasets. This may be correlated with the higher precision of longcallR.

We quantified allele-specific splicing using multiple HG002 datasets from GIAB. On HG002b:MAS:cDNA, a total of 6,227 protein-coding genes and long non-coding RNAs (lncRNAs) are supported by ≥10 phased reads. Out of these genes, longcallR identified 445 genes that contain at least one ASJ (**Supplementary Fig. 1**). 44% of them are also identified with other sequencing technologies applied to the same cell type (**Fig. 3b**) and these shared ASJ genes tend to have more significant P-value: while 47% of ASJ genes shared between MAS-Seq other technologies have P-value < 10^−10^, only 8% of genes specific to MAS-seq have P-value < 10^−10^ (**Supplementary Fig. 2**) – ASJ genes specific to one technology are more prone to statistical fluctuation. We observed more MAS-Seq-specific ASJ genes because this dataset has the highest coverage and thus yields the highest power.

**Fig. 3.**
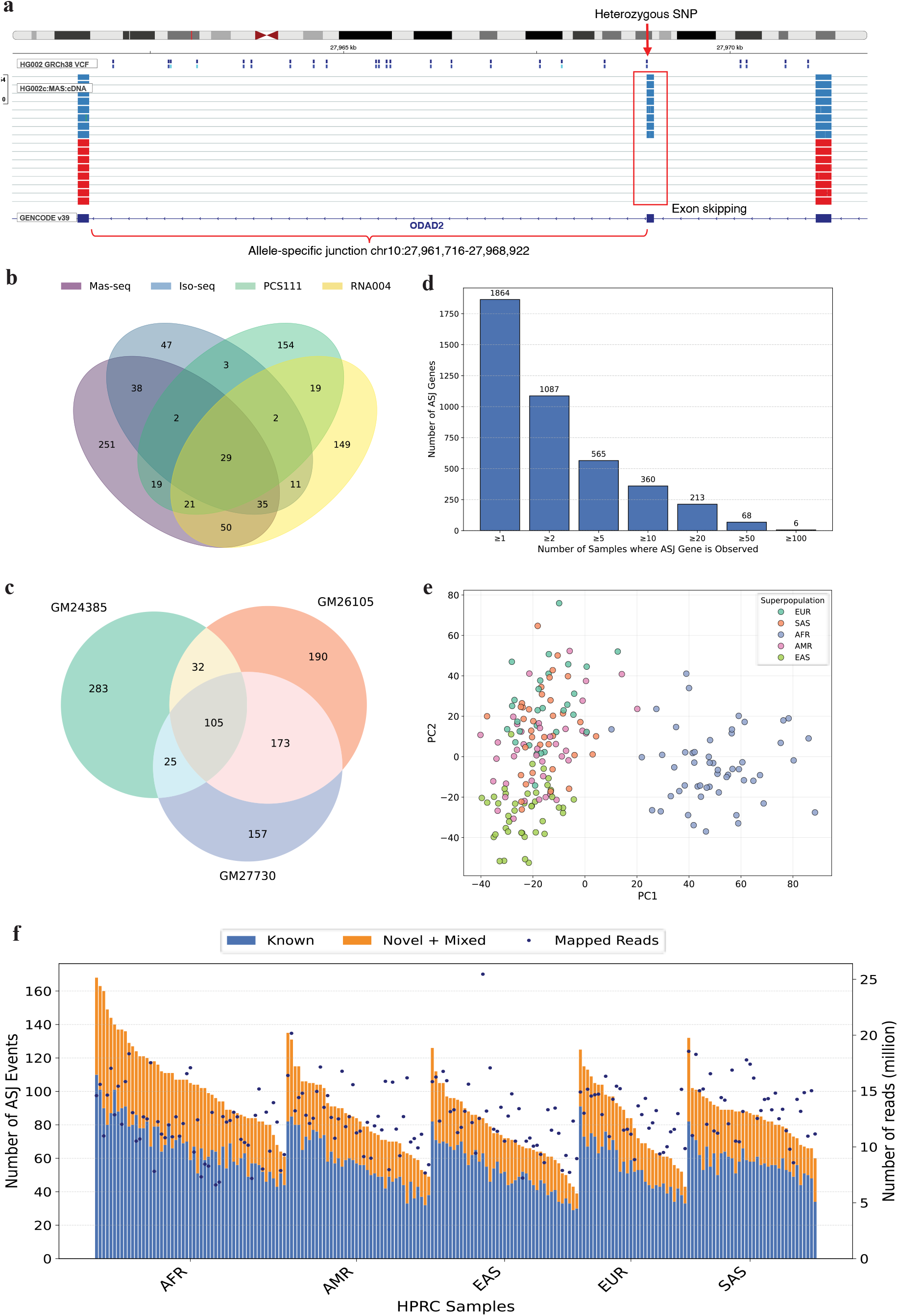
Allele-specific analysis using longcallR. **A**, Example of allele-specific splicing. Reads from haplotype 1 (orange) contain the junction, while reads from haplotype 2 (blue) miss the junction due to an exon-skipping event. A heterozygous SNP at chr10:27,968,923, located 1 bp upstream of the junction. The P-value of this event is 1.0 ×10^−24^. **B**, Comparison of allele-specific junction (ASJ) genes identified in the GM24385 human cell line using PacBio Mas-seq, Iso-seq and Nanopore cDNA (PCS111), direct RNA (RNA004) sequencing. Only protein-coding genes and long non-coding RNAs covered by at least 10 phased reads are analyzed. **C**, Comparison of allele-specific junction (ASJ) genes identified in different cell types GM24385, GM26105 and GM27730 using PacBio Mas-seq. **D**, Cumulative count of significant ASJ genes (P-value < 10^−10^) across HPRC samples. **E**, Principal Component Analysis of ASJ genes across HPRC samples. Each point represents an individual colored by its superpopulation as defined in the 1000 Genomes Project. The input matrix comprises − log_10_ *p* values of ASJ genes that are present in at least five samples at p-value < 0.05. **F**, Count of known and novel ASJ events across HPRC samples. Samples were grouped by superpopulation, with ASJ events classified as purely known (blue) or novel (orange) depending on whether events involve unannotated novel junctions. Dots in dark blue represent the number of mapped reads per sample.

GIAB provides MAS-Seq data for three cell types: GM24385 for EBV-transformed B-lymphocyte, GM26105 for iPSC from B-lymphocyte, and GM27730 for iPSC from fibroblast (**Supplementary Table 1**). Out of 445 ASJ genes identified in GM24385 (dataset H002b:MAS:cDNA), 36% can be found in GM26105 and GM27730 (**Fig. 3d**). GM24385-specific ASJ genes tend to have less significant P-value (**Supplementary Fig. 3**) and have lower read coverage in the other two cell types (**Supplementary Fig. 4**). Having data from multiple cell types helps to increase the power of ASJ gene identification.

Next, we closely investigated the allele-specific junctions in these genes. Note that one alternative splicing event, such as exon skipping, may affect two or more junctions. Therefore, we regard two junctions are related if they share the same donor or acceptor splice sites and we define a junction group, or junction event, to be a set of related junctions via single-linkage clustering such that for each junction, its related junctions must be in the same set (**Fig. 1c**). Among junction groups containing ASJs of P-value < 1 × 10^−10^, 63 contain annotated junctions only, 18 contain novel junctions only and 70 events involve both known and novel junctions (**Supplementary Table 7**, Method). Notably, although only 1.5% of all junctions observed in raw reads are novel, 58.3% of ASJ events contain at least one novel junction – novel junctions are enriched among ASJ events. Furthermore, 46% of significant ASJ events have heterozygous SNPs within a ±10 bp window around the splice sites, compared to only 3% for non-ASJ events.

We applied longcallR to 202 human samples sequenced by the Human Pangenome Reference Consortium (HPRC) with MAS-Seq. We found 1,864 protein-coding and lncRNA genes that contain ASJs with P-value < 10^−10^. 1,087 of them occurred in ≥2 samples (**Fig. 3d**). The principal component analysis (PCA) of ASJ genes suggests the first principal component (PC) separates African and non-African samples while the second PC separates European and East Asian samples (**Fig. 3e**). This result is broadly similar to the PCA analysis of SNPs^20^, which implies that ASJ genes are associated with genetic variants as is expected. On average, each sample has 88 ASJ events identified with P-value < 10^−10^ and 46.5% of the ASJ events contain novel junctions not annotated in GenCode v39 (**Fig. 3f**). Africans have more ASJ events (**Supplementary Fig. 5**), even though they tend to have lower read coverage (**Supplementary Fig. 6**). Within each geographical group, the number of detected ASJ events is correlated with the number of mapped reads (**Fig. 3f**) because we get higher statistical power at higher read coverage.

LongcallR is currently designed for bulk RNA-seq, integrating SNP calling and haplotype phasing at the algorithmic level to achieve highly accurate SNP identification and phasing. Leveraging phased haplotype information, longcallR enables the detection of allele-specific events. We plan to extend longcallR to single-cell RNA-seq, enabling the identification of germline and somatic mutation alongside haplotype phasing at the single-cell level. This extension will allow for a deeper exploration of cell-specific allele-specific genetic effects, providing insights into cellular heterogeneity and regulatory mechanisms.

## Methods

### Overall workflow of longcallR

The longcallR pipeline begins by identifying candidate SNP sites from pileups of reads-to-reference alignment. Next, longcallR-nn constructs a feature image incorporating surrounding sequence and alignment information for each candidate site and applies a neural network to predict genotype and zygosity. Following this, longcallR-phase simultaneously phases SNPs, haplotags reads and filters false-positive calls based on haplotype information. Finally, longcallR utilizes the phased VCF, phased BAM, and annotation to identify allele-specific expression (ASE) and allele-specific junctions (ASJ).

### Identifying candidate SNP sites

LongcallR filters unmapped, secondary, and supplementary reads while applying a mapping quality threshold ≥10. It then counts the occurrences of the four nucleotides (A, C, G, and T) at each position. The depth of a position is determined as the sum of all four nucleotides. The reference allele fraction is calculated as the number of reference alleles divided by the depth, while the alternate allele fraction is computed as the sum of all non-reference alleles divided by the depth. The alternate allele count represents the total number of non-reference alleles. A position is selected as a candidate SNP if it meets the following criteria: a sequencing depth ≥6, an alternate allele fraction ≥0.1, and an alternate allele count ≥2.

### longcallR-nn

*Model architecture*. longcallR-nn is a deep-learning-based SNP caller that leverages ResNet to analyze pileup images generated from read-to-reference alignments. The process of generating a pileup image begins by extracting multiple feature layers from sequencing data, including Base, Reference, Insertion, Base Quality, Mapping Quality, Read Strand, and Transcript Strand. The Base layer encodes the nucleotide bases observed at the position, while the Reference layer represents the reference genome sequence. The Insertion layer indicates the presence of insertions at the site. The Base Quality layer provides quality scores of individual bases, while the Mapping Quality layer represents the alignment score of each read. The Read Strand layer specifies the strand orientation of each read, whereas the Transcript Strand layer represents the inferred transcript strand for the read by aligner. These feature layers are stacked together to form a multi-channel representation of the candidate SNP site. The resulting pileup image has dimensions of 41 × *d* × 7, where 41 represents the window size surrounding the SNP site, *d* denotes the number of reads covering the site, and 7 corresponds to the number of feature channels. Since some feature layers have different sizes, padding is applied to ensure uniform dimensions. In the Reference layer, the reference sequence is duplicated *d* times to form a 41 × *d* matrix. In the Mapping Quality layer, as each read has a single mapping quality value, this value is replicated 41 times to match the 41 × *d* format. Similarly, in the Read Strand layer and Transcript Strand layer, where each read has a single strand value, the value is repeated 41 times to conform to the 41 × *d* dimensions. The pileup image is then fed into a convolution neural network, ResNet-50, which learns the relationships between genotype and sequence context. The model output is subsequently used for two classification tasks: zygosity prediction and genotype prediction. Zygosity classification includes five categories: homozygous reference (0/0), heterozygous variant (0/1), homozygous variant (1/1), biallelic variant (1/2), and RNA editing. Genotype classification determines combinations of reference alleles {A, C, G, T} and alternate alleles {A, C, G, T}. The model parameters are optimized using stochastic gradient descent (SGD) with an initial learning rate of 0.01.

*Training dataset*. Raw sequence reads are aligned to the GRCh38 reference genome using the minimap2 splice mode, followed by read coverage calculation at each genomic position using samtools. Candidate SNP sites are selected based on a minimum coverage threshold ≥6 and an alternative allele fraction ≥0.1, while those located outside high-confidence regions are filtered out. For labeling, each candidate is checked against REDIportal^21^, a comprehensive database of A-to-I RNA editing events. If the candidate’s position matches a record in REDIportal, the site was labeled as an A-to-I RNA editing event. For candidates without a match, the position was compared against the ground truth DNA SNP set to determine its label. We used sample WTC11 from LRGASP (Long-read RNA-seq Genome Annotation Assessment Project)^22^ to train the Nanopore cDNA model. Genomic variants produced by the Allen Institute were taken as the ground truth. The rest of the models were trained on HG002 samples with the genomic variants from GIAB as the ground truth. We used Kinnex full-length RNA dataset (HG002a:MAS:cDNA) from PacBio and Mas-Seq dataset (HG002b:MAS:cDNA) from GIAB for training the PacBio Mas-Seq model, used GIAB Iso-Seq data produced by Baylor College of Medicine for the PacBio Iso-Seq model, and used Nanopore dRNA dataset generated by Clair3-RNA developers for the dRNA model (**Supplementary Table 1**).

### longcallR-phase

*Algorithmic framework*. longcallR-phase is an algorithm designed to simultaneously SNP calling and phasing for long-read RNA-seq data. The core concept of the algorithm is structured as follows. Initially, a subset of high-confidence true heterozygous SNPs is selected from all candidate SNPs. longcallR-phase then phases this subset of SNPs and assigns haplotype tags to the reads. Subsequently, by leveraging the read haplotype information, the algorithm validates remaining candidates and identifies more compatible SNPs to expand the subset used for phasing. As the number of phased SNPs increases, read haplotype tagging is further refined and improved. By iteratively repeating this process, the algorithm effectively distinguishes true SNPs from sequencing errors and RNA editing events.

Suppose there are *n* fragments and *m* heterozygous SNPs. Let *I*_*k*_ denote the set of SNPs present on the fragment *k*, and similarly, let *K*_*i*_ represent the set of fragments overlapping with SNP *i*. Define *δ*_*i*_ = {1, −1} as the haplotype of SNP *i* and *σ*_*k*_ = {1, −1} as the haplotag of fragment *k*. At each SNP, the reference allele is assigned to value 1 and the alternate allele to value −1. The probability of observing allele *x* (*x* ∈ {1, −1}) at SNP *i* on fragment *k* is denoted by *q*_*ki*_(*x*), which is related to the base quality of allele at SNP *i* on fragment *k*. The objective is to find a configuration {*δ, σ*} that maximizes the overall probability:

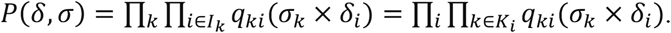

Initially, *δ* and *σ* are assigned randomly. The process then involves two steps:

1. Fix *δ* and update *σ*_*k*_: flip each *σ*_*k*_ sequentially and retain the flip if it increases the conditional probability:

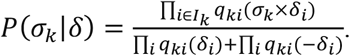
2. Fix *σ* and update *δ*_*i*_: flip each *δ*_*i*_ sequentially and retain the flip if it increases the conditional probability:

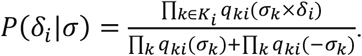

These steps are repeated iteratively, alternating between updating *δ* and *σ*, until the overall probability *P*(*δ, σ*) converges and no longer increases. Finally, based on the haplotype tagging of reads, the confidence of candidate SNP is quantified using a score calculated as −10 log_10_(*P*(*δ*_*i*_|*σ*)). If the score exceeds a predefined threshold, the candidate SNP is identified as a true germline heterozygous SNP. Meanwhile, each read is assigned to one haplotype by comparing the probabilities *P*(*σ*_*k*_|*δ*) and *P*(−*σ*_*k*_|*δ*).

*Heuristic initialization*. The final phasing is obtained by iteratively adjusting the phase of individual SNPs and their corresponding reads. However, the presence of false-positive SNPs or RNA-editing events among candidate SNPs may introduce unnecessary phase flips during this process. To enhance algorithmic convergence, our approach first identifies a subset of reliable SNPs, establishes a more reasonable phasing initialization compared to a fully random initialization, and then gradually incorporates more challenging SNPs into the process. Let *S*_*i*_ and *S*_*j*_ denote any two SNPs in the candidate set, where *A* and *a* represent two allele types at *S*_*i*_, and *B* and *b* represent two allele types at *S*_*j*_. For all possible allele combinations {(*A, B*), (*A, b*), (*a, B*), (*a, b*)}, we count the number of reads covering both *S*_*i*_ and *S*_*j*_ that correspond to each combination. Using these counts, we compute:

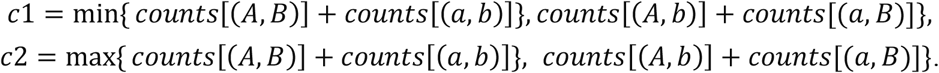

The score is then defined as *c*1/*c*2, where a score of 0 indicates that *S*_*i*_ and *S*_*j*_ can be phased into the same haplotype without conflicts. We construct a graph representing all SNP pairs in the candidate set. For each pair with a score of 0, *S*_*i*_ and *S*_*j*_ are treated as two nodes in a graph, with an edge added between them. After processing all SNP pairs, connected components within the graph are identified. SNPs within the same connected component are inferred to belong to the same haplotype and are initialized with the identical phase. During subsequent iterations, these SNPs are either flipped simultaneously or remain unchanged as a group.

### Allele-specific analysis

*Allele-specific expression identification*. To assign reads to each gene, we first extract exon regions from the provided annotation. Since individual reads may span multiple genes, we parse the CIGAR string of each read alignment to calculate the number of mapped bases within the exon regions. Each read is then assigned to the gene with which it has the highest number of mapped exon bases. Subsequently, for each gene, the assigned reads are collected and integrated with read haplotype information to determine the read counts corresponding to each haplotype. If multiple phase sets are found among the reads, the phase set containing the highest number of reads is selected, and the haplotype-specific read counts are computed within that phase set. Finally, a beta-binomial test is performed to assess whether the observed counts of reads for the two haplotypes were significantly different from the expected ratio of 0.5 and the p-values of allele-specific expression were corrected for multiple hypothesis testing with the Benjamini–Hochberg (BH) procedure (FDR<5%). When paternal and maternal information for all DNA SNPs is available, we take the intersection of phased RNA SNPs and phased DNA SNPs. The paternal and maternal haplotypes are then determined by comparing the number of paternal and maternal alleles among the overlapping SNPs in the gene. This allows allele-specific expression (ASE) to be further classified into paternal or maternal ASE.

*Allele-specific junction identification*. To identify allele-specific junctions, we first generate a set of candidate junctions by extracting all junctions with canonical GT-AG splice site signals from raw mapped reads, followed by merging those with identical start and end coordinates. For a given query junction within the candidate set, we classify all overlapping reads into two groups: those where the junction is present and those where it is absent. The junctions without sufficient overlapped reads (10x) are filtered out. Reads containing a junction with identical start and end positions as the query junction are classified as present reads (*R*_4_), while all other reads are classified as absent reads (*R*_*a*_). Haplotype phasing information is then utilized to further partition these reads into haplotype-specific groups. Present reads are divided into two sets: *haplotype 1 present* (*R*_41_) and *haplotype 2 present* (*R*_4*2*_). Similarly, absent reads are divided into *haplotype 1 absent* (*R*_*a*1_) and *haplotype 2 absent* (*R*_*a2*_). If multiple phase sets are found among the reads, only the phase set with the highest number of reads is retained, and reads from other phase sets are excluded from the analysis. When genomic SNPs are available, reads phased exclusively by RNA SNPs but not overlapping any genomic SNPs, which may arise from technical artifacts, are additionally filtered. Finally, a Fisher’s exact test is applied to evaluate the dependency between haplotype and junction presence using a 2 × 2 contingency table based on the sizes of the four groups (*R*_*p*1_, *R*_*p*2_, *R*_*a*1_, *R*_*a*2_), thereby identifying junctions that exhibit allele-specific behavior. The p-values of allele-specific junctions were corrected for multiple hypothesis testing using the Benjamini–Hochberg (BH) procedure (FDR<5%). Additionally, the symmetric odds ratio (SOR) is calculated using these four values as follows:

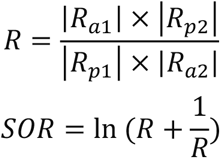

The read coverage, SOR and number of reads from both haplotypes (|*R*_*a*1_| + Y*R*_41_Y, |*R*_*a2*_| + Y*R*_4*2*_Y) are key factors influencing the statistical power of detection. To simplify the power analysis, we assume that the number of reads from both haplotypes is identical. We then assess the power to detect allele-specific junctions under varying read coverages and SOR through simulations. For a given coverage, we use two multinomial distributions to randomly generate the number of reads absent (|*R*_*ai*_|) and present (Y*R*_4*i*_Y) at the junction in each haplotype, reflecting the specified SOR of interest. Significant ASJs are identified using Fisher’s exact test (p-value < 0.01). Our results show that for an SOR effect size of 2.0, the statistical power exceeds 80% when the total read coverage is greater than 50×. Since a single alternative splicing event may involve two or more junctions, we group related junctions into a single junction event if they share either a start or end position. For each junction event, if at least one junction has a P-value < 0.05, it is classified as an ASJ event; otherwise, it is considered a non-ASJ event. ASJ events are further categorized as significant or insignificant. An ASJ event is deemed significant if it contains at least one junction with a P-value < 10^−10^ and SOR ≥ 2; otherwise, it is considered insignificant. Each ASJ event is additionally classified as purely known, purely novel, or mixed. If all junctions within the ASJ event with P-value < 0.05 are annotated as known, the event is classified as purely known. If all such junctions are novel, it is classified as purely novel. Events containing both known and novel junctions are classified as mixed ASJ events.

### Method Benchmarking

*Benchmarking of SNP calling performance*. To generate reliable benchmarks for each dataset, we first calculated read coverage using samtools to obtain well-expressed regions with coverage ≥20. These expressed regions were then intersected with the high-confidence regions from GIAB to define reliable benchmarking regions. Finally, DNA SNPs sourced from GIAB within these benchmarking regions were taken as the ground truth RNA SNPs for each lrRNA-seq dataset. Using these ground truth RNA SNPs, we assessed precision and recall for SNP calls with *hap*.*py*^25^.

*Benchmarking of haplotype phasing performance*. Phasing performance is assessed using several metrics, including total switch errors, average switch error rate, total blockwise Hamming errors, average blockwise Hamming error rate, and phase block N50. Total switch errors and total blockwise Hamming errors represent the sum of errors across all chromosomes, while the average switch error rate and average blockwise Hamming error rate denote the mean error rate across different chromosomes. These metrics are computed for chromosomes 1–22. The phased VCFs generated by longcallR and WhatsHap are compared with the GIAB phased VCF calls using the WhatsHap *compare* module to determine switch and Hamming errors. The WhatsHap *stats* module is used to generate GTF files of SNV phase blocks, which are subsequently utilized to calculate the phase block N50.

## Data Availability

The raw reads for HG002a:MAS:cDNA are available at https://downloads.pacbcloud.com/public/dataset/Kinnex-full-length-RNA/DATA-Revio-HG002-1/. The raw reads for HG002b:MAS:cDNA, HG002c:MAS:cDNA, HG002d:MAS:cDNA, HG004:MAS:cDNA, HG005:MAS:cDNA, HG002a:ISO:cDNA, HG002b:ISO:cDNA, HG002c:ISO:cDNA, HG004:ISO:cDNA, HG005:ISO:cDNA can be accessed at https://ftp-trace.ncbi.nlm.nih.gov/giab/ftp/data_RNAseq/. The WTC11:ONT:cDNA reads can be found in ENCODE under ENCSR539ZXJ. The raw reads for HG002a:ONT:cDNA, HG002b:ONT:cDNA, HG004:ONT:cDNA and HG005:ONT:cDNA are available at https://s3.amazonaws.com/gtl-publicdata/giab/bams/cDNA/. The HG002:ONT:dRNA, HG004:ONT:dRNA, and HG005:ONT:dRNA datasets, generated by the University of Hong Kong, are available in NCBI under SRX26304755, SRX26304756, SRX26304757. HPRC human samples are available at s3://human-pangenomics/working/HPRC/. For ground truth DNA variants, the VCF files for HG002, HG004, and HG005 are available at GIAB v4.2.1. The VCF for WTC11 is provided by the Allen Institute at https://open.quiltdata.com/b/allencell/tree/aics/wtc11_short_read_genome_sequence/. The reference genome GRCh38 can be downloaded from NCBI.

## Code Availability

LongcallR is available at https://github.com/huangnengCSU/longcallR. LongcallR-nn is available at https://github.com/huangnengCSU/longcallR-nn. LongcallR scripts are available at https://github.com/huangnengCSU/longcallR_scripts. The Nextflow of longcallR is available at https://github.com/huangnengCSU/longcallR-nf.

## Acknowledgement

We would like to acknowledge the National Genome Research Institute (NHGRI) for funding the following grants supporting the creation of the human pangenome reference: U41HG010972, U01HG010971, U01HG013760, U01HG013755, U01HG013748, U01HG013744, R01HG011274, and the Human Pangenome

Reference Consortium (BioProject ID: PRJNA730823).

## Supplementary Tables

**Supplementary Table 1.**
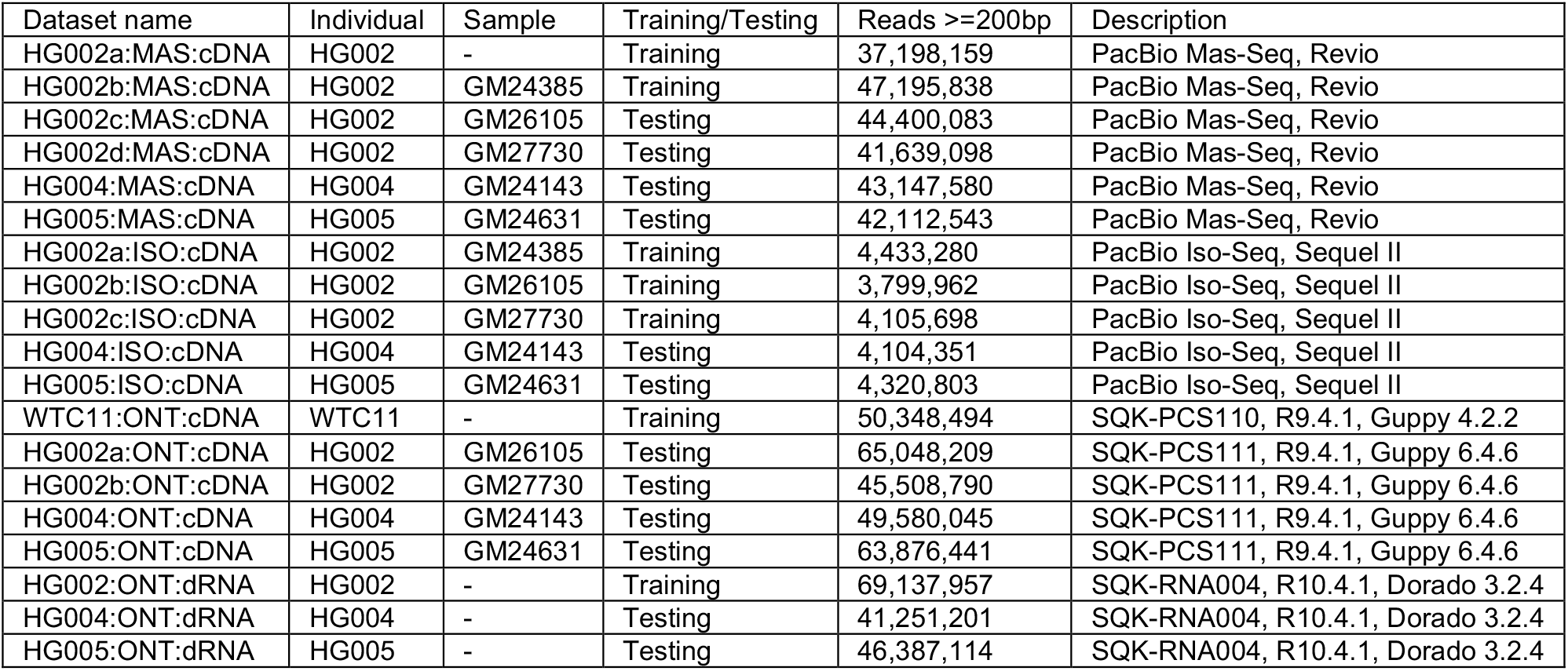
Summary of datasets.

**Supplementary Table 2.** Evaluation of SNP calls by longcallR and Clair3-RNA across whole-genome regions.

|  | Recall |  | Precision |  | F1 score |  | TP |  |
| --- | --- | --- | --- | --- | --- | --- | --- | --- |
|  | longcallR | Clair3-RNA | longcallR | Clair3-RNA | longcallR | Clair3-RNA | longcallR | Clair3-RNA |
| HG002c:MAS:cDNA | 94.90% | 94.82% | 98.56% | 97.64% | 96.70% | 96.21% | 93,320 | 93,244 |
| HG002d:MAS:cDNA | 94.76% | 94.79% | 98.65% | 97.79% | 96.67% | 96.27% | 96,874 | 96,905 |
| HG004:MAS:cDNA | 93.85% | 94.12% | 98.53% | 97.60% | 96.13% | 95.83% | 90,073 | 90,332 |
| HG005:MAS:cDNA | 95.28% | 95.59% | 98.78% | 97.81% | 97.00% | 96.69% | 111,289 | 111,654 |
| HG004:ISO:cDNA | 91.62% | 94.78% | 98.75% | 97.62% | 95.05% | 96.18% | 34,373 | 35,559 |
| HG005:ISO:cDNA | 90.98% | 93.92% | 99.03% | 97.58% | 94.84% | 95.72% | 34,167 | 35,269 |
| HG002a:ONT:cDNA | 73.90% | 81.71% | 85.83% | 72.51% | 79.42% | 76.83% | 26,939 | 29,783 |
| HG002b:ONT:cDNA | 77.91% | 85.06% | 88.46% | 78.86% | 82.85% | 81.85% | 27,601 | 30,135 |
| HG004:ONT:cDNA | 78.04% | 83.28% | 91.68% | 85.92% | 84.31% | 84.58% | 20,126 | 21,477 |
| HG005:ONT:cDNA | 82.45% | 86.74% | 93.88% | 89.44% | 87.79% | 88.07% | 28,296 | 29,769 |
| HG004:ONT:dRNA | 85.82% | 93.78% | 97.95% | 91.26% | 91.48% | 92.51% | 38,703 | 42,297 |
| HG005:ONT:dRNA | 87.34% | 94.96% | 98.04% | 91.15% | 92.38% | 93.02% | 41,665 | 45,296 |

**Supplementary Table 3.** Evaluation of SNP calls by longcallR and Clair3-RNA in exon regions.

|  | Recall |  | Precision |  | F1 score |  | TP |  |
| --- | --- | --- | --- | --- | --- | --- | --- | --- |
|  | longcallR | Clair3-RNA | longcallR | Clair3-RNA | longcallR | Clair3-RNA | longcallR | Clair3-RNA |
| HG002c:MAS:cDNA | 95.43% | 94.01% | 98.94% | 98.08% | 97.16% | 96.00% | 45,002 | 44,334 |
| HG002d:MAS:cDNA | 95.24% | 93.93% | 98.93% | 98.08% | 97.05% | 95.96% | 44,464 | 43,849 |
| HG004:MAS:cDNA | 93.10% | 92.26% | 98.72% | 97.76% | 95.83% | 94.93% | 38,116 | 37,770 |
| HG005:MAS:cDNA | 94.40% | 93.48% | 98.86% | 97.87% | 96.58% | 95.62% | 39,988 | 39,597 |
| HG004:ISO:cDNA | 90.61% | 94.33% | 99.05% | 98.05% | 94.64% | 96.15% | 20,347 | 21,183 |
| HG005:ISO:cDNA | 90.89% | 93.66% | 99.14% | 97.89% | 94.84% | 95.73% | 20,618 | 21,246 |
| HG002a:ONT:cDNA | 77.74% | 84.11% | 86.72% | 72.77% | 81.99% | 78.03% | 22,719 | 24,578 |
| HG002b:ONT:cDNA | 81.52% | 87.41% | 89.28% | 79.99% | 85.22% | 83.53% | 23,418 | 25,109 |
| HG004:ONT:cDNA | 81.15% | 85.71% | 93.85% | 89.26% | 87.04% | 87.45% | 17,826 | 18,827 |
| HG005:ONT:cDNA | 85.30% | 88.76% | 95.55% | 91.99% | 90.13% | 90.35% | 23,417 | 24,368 |
| HG004:ONT:dRNA | 86.81% | 94.85% | 98.50% | 93.06% | 92.29% | 93.95% | 30,472 | 33,294 |
| HG005:ONT:dRNA | 88.09% | 95.83% | 98.54% | 92.71% | 93.02% | 94.24% | 31,164 | 33,903 |

**Supplementary Table 4.** Evaluation of variant calling by only longcallR-nn (nn-only), longcallR-nn plus longcallR-phase (nn + phase) and only longcallR-phase (phase-only).

|  | Methods | HG004:<br>MAS:<br>cDNA | HG005:<br>MAS:<br>cDNA | HG004:<br>ISO:<br>cDNA | HG005:<br>ISO:<br>cDNA | HG004:<br>ONT:<br>cDNA | HG005:<br>ONT:<br>cDNA | HG004:<br>ONT:<br>dRNA | HG005:<br>ONT:<br>dRNA |
| --- | --- | --- | --- | --- | --- | --- | --- | --- | --- |
| Recall | nn-only | 94.81% | 95.90% | 93.36% | 92.57% | 83.44% | 86.51% | 90.28% | 91.34% |
|  | nn + phase | 93.85% | 95.28% | 91.62% | 90.98% | 78.04% | 82.45% | 85.82% | 87.34% |
|  | phase-only | 92.06% | 93.09% | 91.07% | 88.86% | 65.38% | 64.05% | 63.10% | 62.83% |
| Precision | nn-only | 97.95% | 98.16% | 98.32% | 98.32% | 86.42% | 87.19% | 97.18% | 97.10% |
|  | nn + phase | 98.53% | 98.78% | 98.75% | 99.03% | 91.68% | 93.88% | 97.95% | 98.04% |
|  | phase-only | 98.18% | 98.44% | 98.55% | 98.69% | 92.51% | 95.65% | 96.74% | 97.14% |

**Supplementary Table 5.**
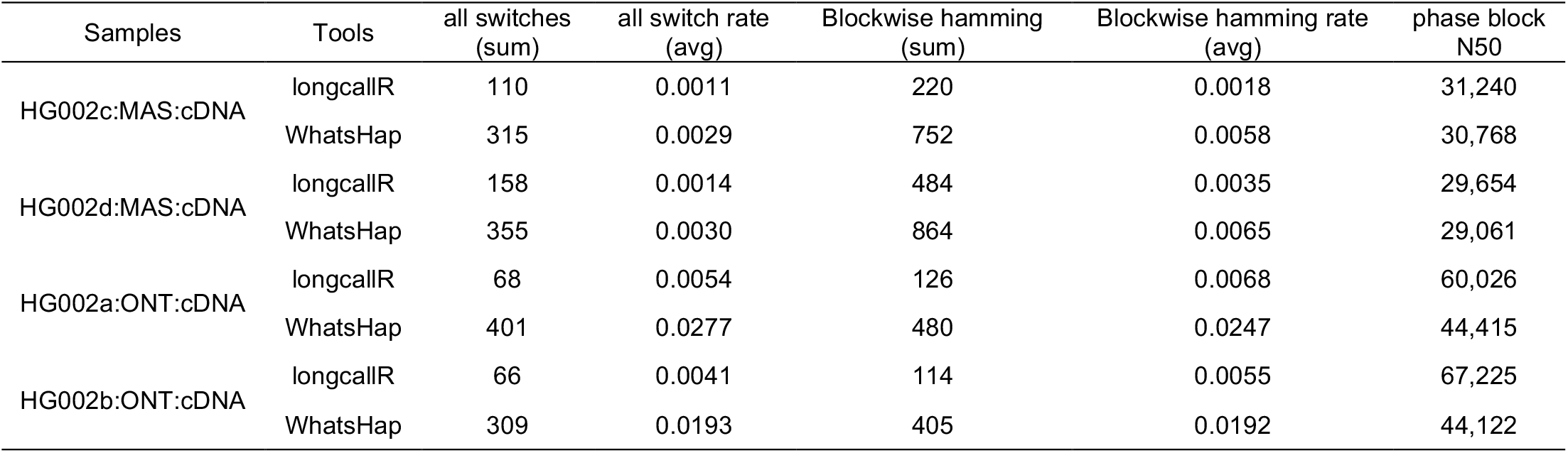
Evaluation of phasing results from longcallR-phase and WhatsHap for SNPs identified by longcallR-nn.

| Samples | Tools | all switches<br>(sum) | all switch rate<br>(avg) | Blockwise hamming<br>(sum) | Blockwise hamming rate<br>(avg) | phase block<br>N50 |
| --- | --- | --- | --- | --- | --- | --- |
| HG002c:MAS:cDNA | longcallR | 110 | 0.0011 | 220 | 0.0018 | 31,240 |
|  | WhatsHap | 315 | 0.0029 | 752 | 0.0058 | 30,768 |
| HG002d:MAS:cDNA | longcallR | 158 | 0.0014 | 484 | 0.0035 | 29,654 |
|  | WhatsHap | 355 | 0.0030 | 864 | 0.0065 | 29,061 |
| HG002a:ONT:cDNA | longcallR | 68 | 0.0054 | 126 | 0.0068 | 60,026 |
|  | WhatsHap | 401 | 0.0277 | 480 | 0.0247 | 44,415 |
| HG002b:ONT:cDNA | longcallR | 66 | 0.0041 | 114 | 0.0055 | 67,225 |
|  | WhatsHap | 309 | 0.0193 | 405 | 0.0192 | 44,122 |

**Supplementary Table 6.** Evaluation of phasing results from longcallR-phase and WhatsHap for SNPs identified by Clair3-RNA.

| Samples | Tools | all switches<br>(sum) | all switch rate<br>(avg) | Blockwise hamming<br>(sum) | Blockwise hamming rate<br>(avg) | phase block<br>N50 |
| --- | --- | --- | --- | --- | --- | --- |
| HG002c:MAS:cDNA | longcallR | 147 | 0.0012 | 375 | 0.0025 | 26,207 |
|  | WhatsHap | 290 | 0.0024 | 721 | 0.0047 | 27,488 |
| HG002d:MAS:cDNA | longcallR | 160 | 0.0012 | 471 | 0.0030 | 24,569 |
|  | WhatsHap | 309 | 0.0024 | 805 | 0.0054 | 26,263 |
| HG002a:ONT:cDNA | longcallR | 88 | 0.0056 | 135 | 0.0060 | 52,051 |
|  | WhatsHap | 210 | 0.0120 | 314 | 0.0130 | 40,367 |
| HG002b:ONT:cDNA | longcallR | 104 | 0.0061 | 198 | 0.0086 | 57,016 |
|  | WhatsHap | 217 | 0.0106 | 318 | 0.0114 | 40,865 |

**Supplementary Table 7.** Summary of ASJ events in the human cell line GM24385 using different sequencing technologies. ASJ events are categorized as entirely known, entirely novel, or mixed based on existing transcript annotations. ASJ events with a nearby SNP are defined as those in which a DNA SNP is located within +/-10bp of the junction.

| Dataset | Category of ASJ events | Number of ASJ events | ASJ events with nearby SNP |
| --- | --- | --- | --- |
| GM24385<br>PacBio Mas-seq | Entirely known | 63 | 29 |
|  | Entirely novel | 18 | 7 |
|  | Mixed | 70 | 34 |
| GM24385<br>PacBio Iso-seq | Entirely known | 34 | 12 |
|  | Entirely novel | 2 | 1 |
|  | Mixed | 22 | 9 |
| GM24385<br>ONT PCS111 | Entirely known | 13 | 5 |
|  | Entirely novel | 4 | 3 |
|  | Mixed | 17 | 9 |
| GM24385<br>ONT RNA004 | Entirely known | 42 | 19 |
|  | Entirely novel | 15 | 5 |
|  | Mixed | 59 | 30 |

## Supplementary Figures

**Supplementary Figure 1.**
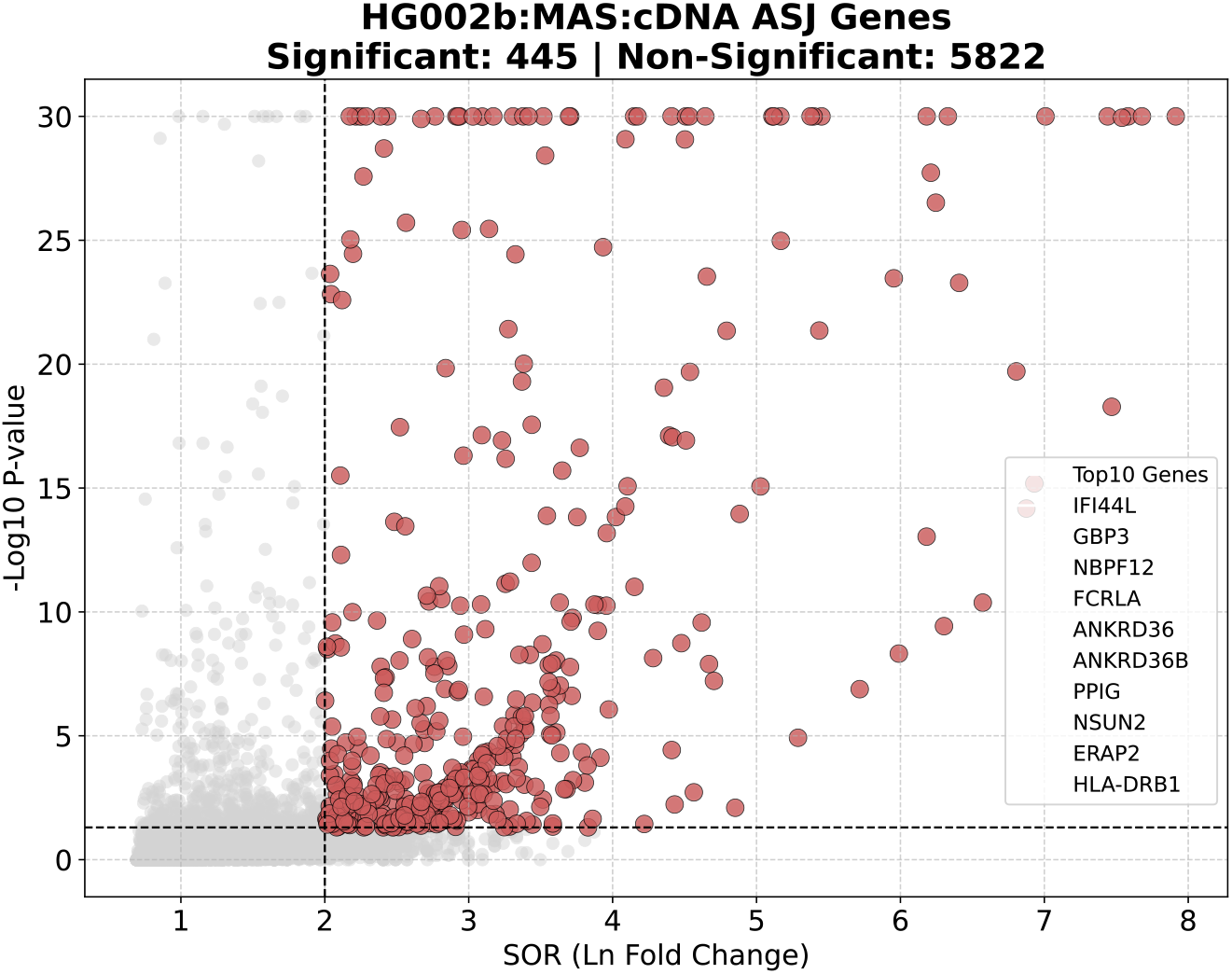
Volcano-like plot illustrating genes with allele-specific junctions (ASJs) identified in the HG002b:MAS:cDNA dataset using longcallR. The x-axis represents the symmetric odds ratio (SOR; Methods), while the y-axis represents the -log_10_ p-value. Each point corresponds to one gene, with significant ASJ genes highlighted in red and non-significant genes shown in gray. The vertical dashed line indicates the SOR threshold of 2, and the horizontal dashed line represents the p-value threshold of 0.05. The top 10 ASJ genes with the lowest p-values are listed in the legend.

**Supplementary Figure 2.**
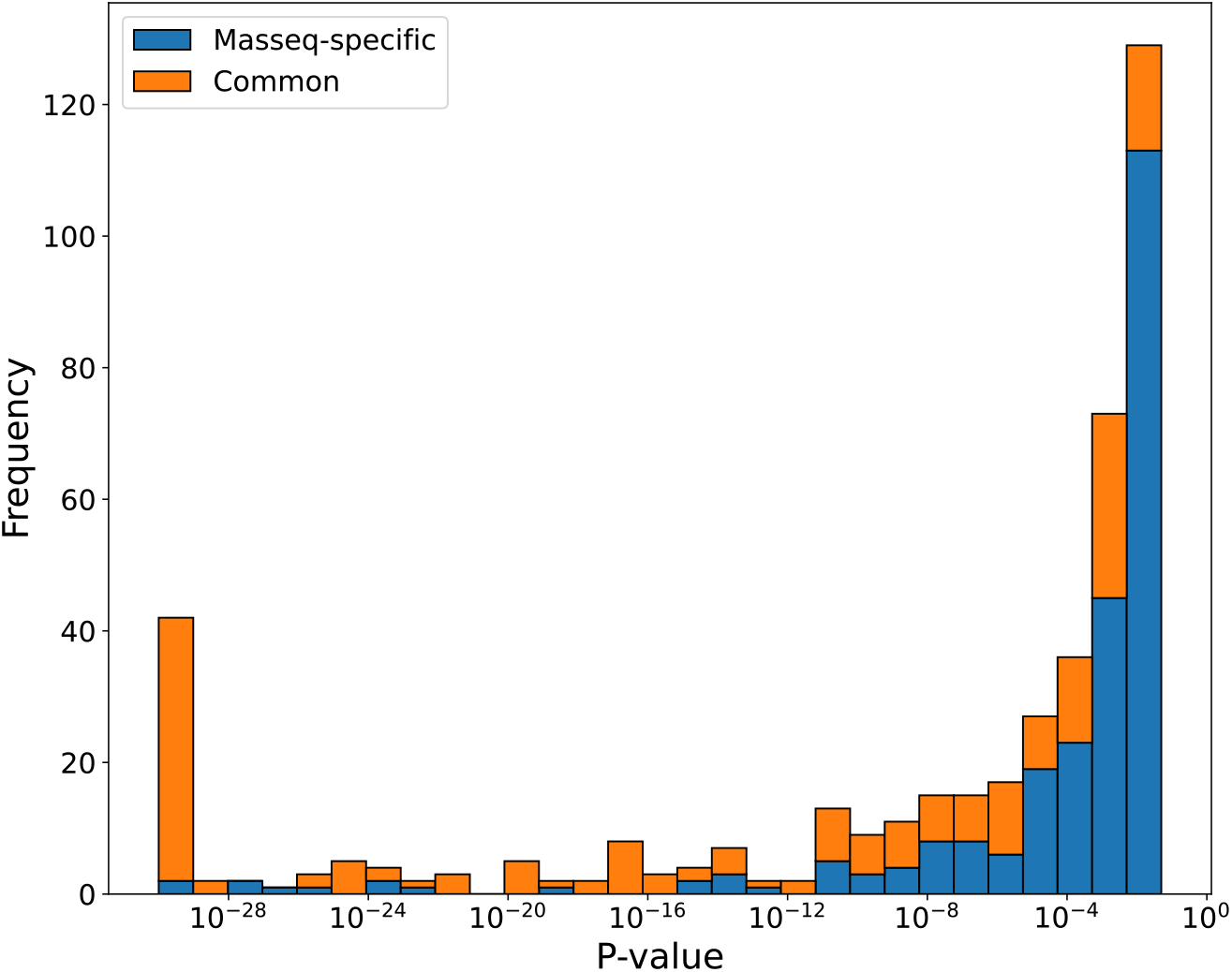
Distribution of p-values for ASJ genes that are either specific to PacBio Mas-seq or shared with at least one other sequencing technology (PacBio Iso-seq, Nanopore cDNA, or Nanopore direct RNA) in the GM24385 human cell line. Blue bars represent ASJ genes detected exclusively by Mas-seq, while orange bars indicate ASJ genes identified by Mas-seq and at least one additional sequencing platform.

**Supplementary Figure 3.**
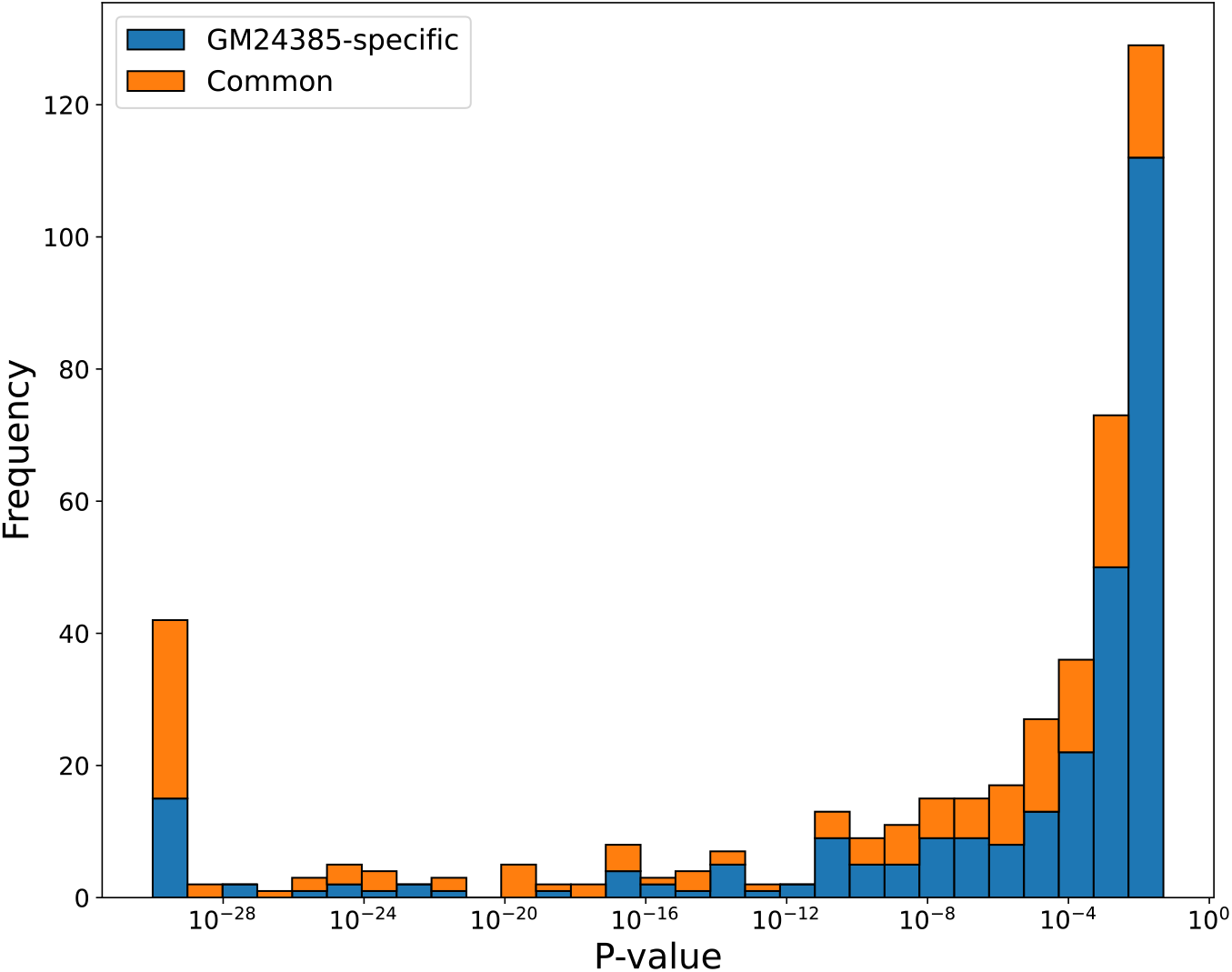
Distribution of p-values for ASJ genes that are either specific to the GM24385 cell line or shared with at least one additional cell line (GM26105 or GM27730), based on PacBio Mas-seq data. Blue bars represent ASJ genes detected exclusively in GM24385, while orange bars indicate ASJ genes identified in GM24385 and at least one other cell line.

**Supplementary Figure 4.**
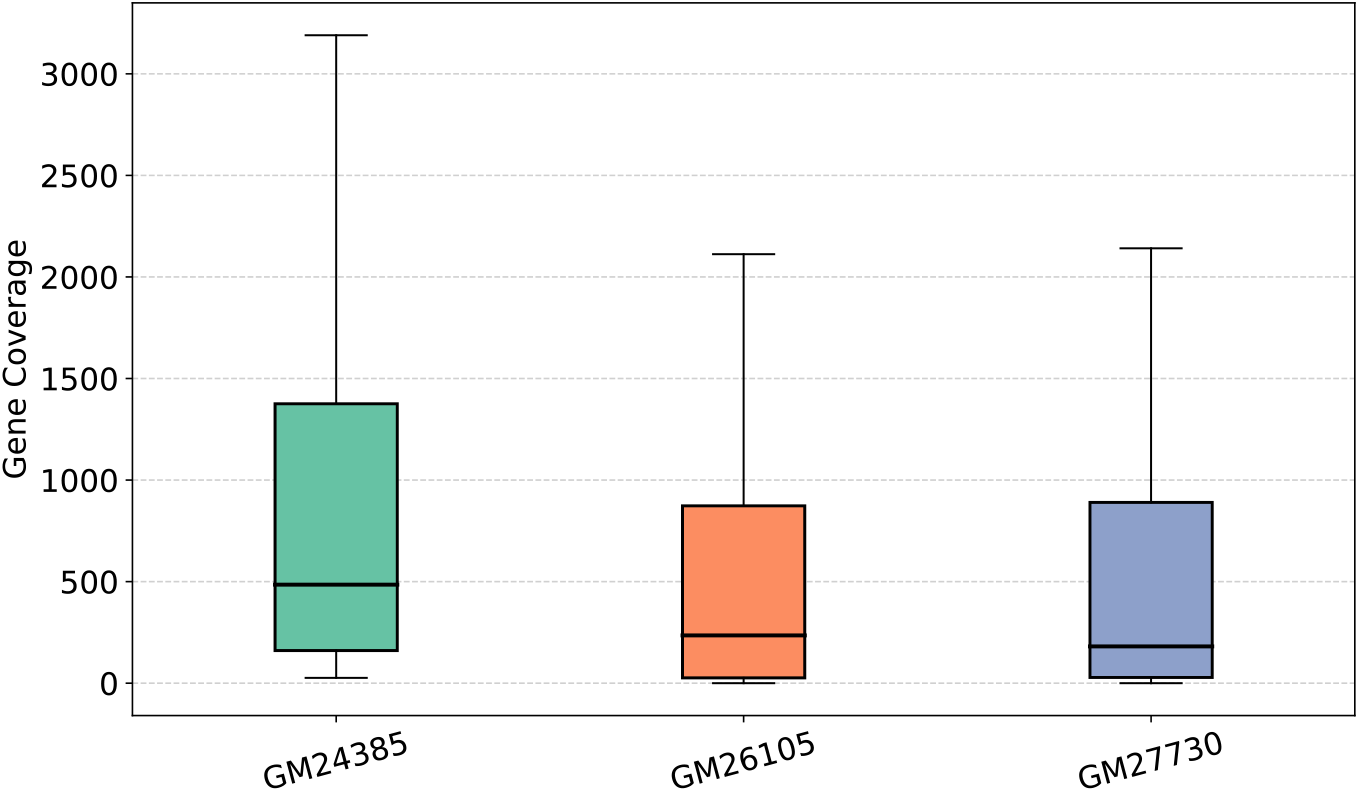
Box plot of gene coverage for GM24385-specific ASJ genes across three cell lines: GM24385, GM26105, and GM27730. Gene coverage is significantly higher in GM24385 compared to GM26105 and GM27730, which show comparable coverages.

**Supplementary Figure 5.**
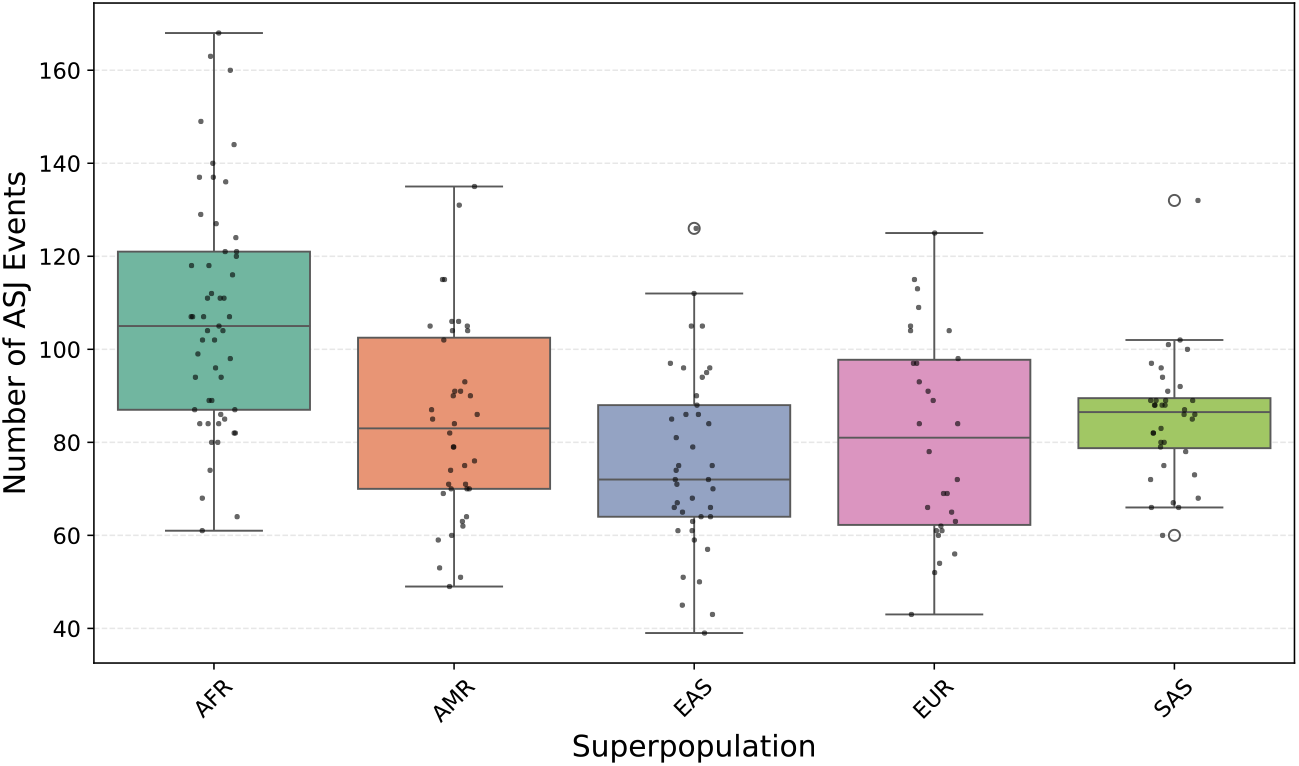
Box plot of ASJ events across HPRC samples, grouped by superpopulation. Each dot represents one sample. Among the five groups, AFR shows the highest number of ASJ events, while EAS displays the lowest. AMR, EUR, and SAS exhibit comparable ASJ event counts.

**Supplementary Figure 6.**
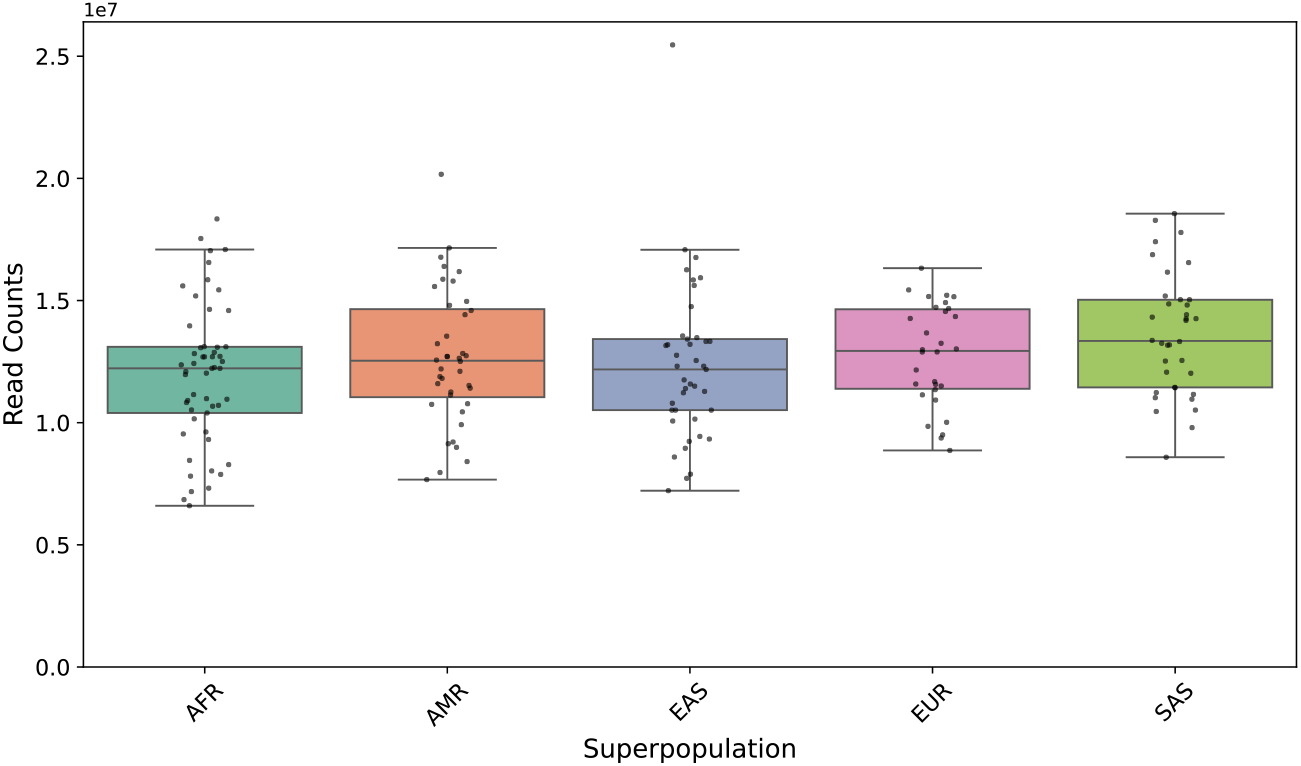
Box plot of mapped read counts across HPRC samples, grouped by superpopulation. Each dot represents one individual sample. AFR and EAS samples show relatively lower read counts, while AMR, EUR, and SAS samples exhibit higher and comparable read counts.

## Supplementary Notes

Supplementary Note 1. Software commands

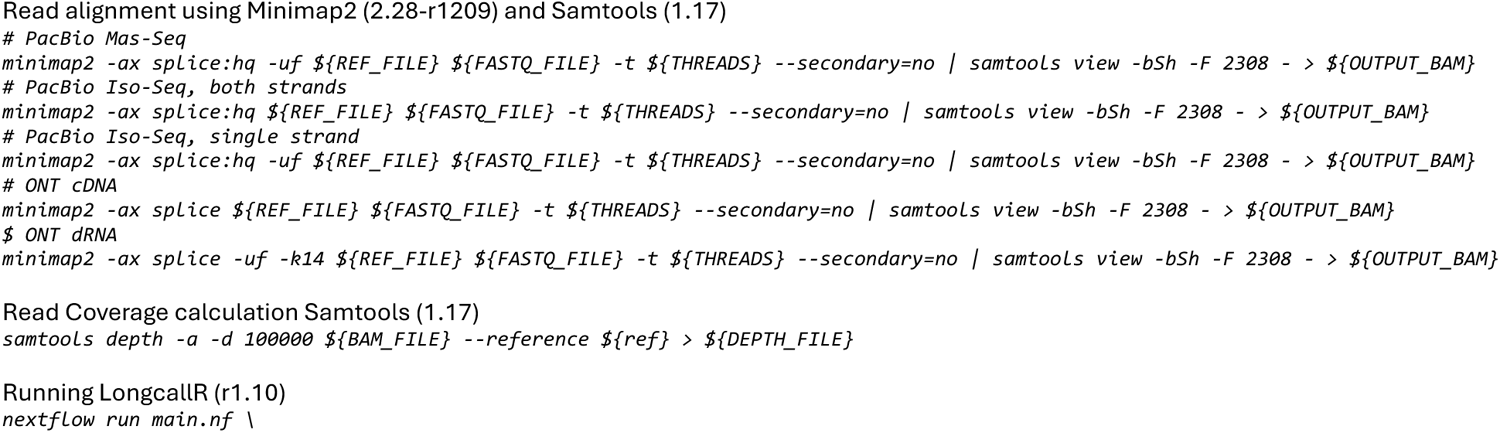
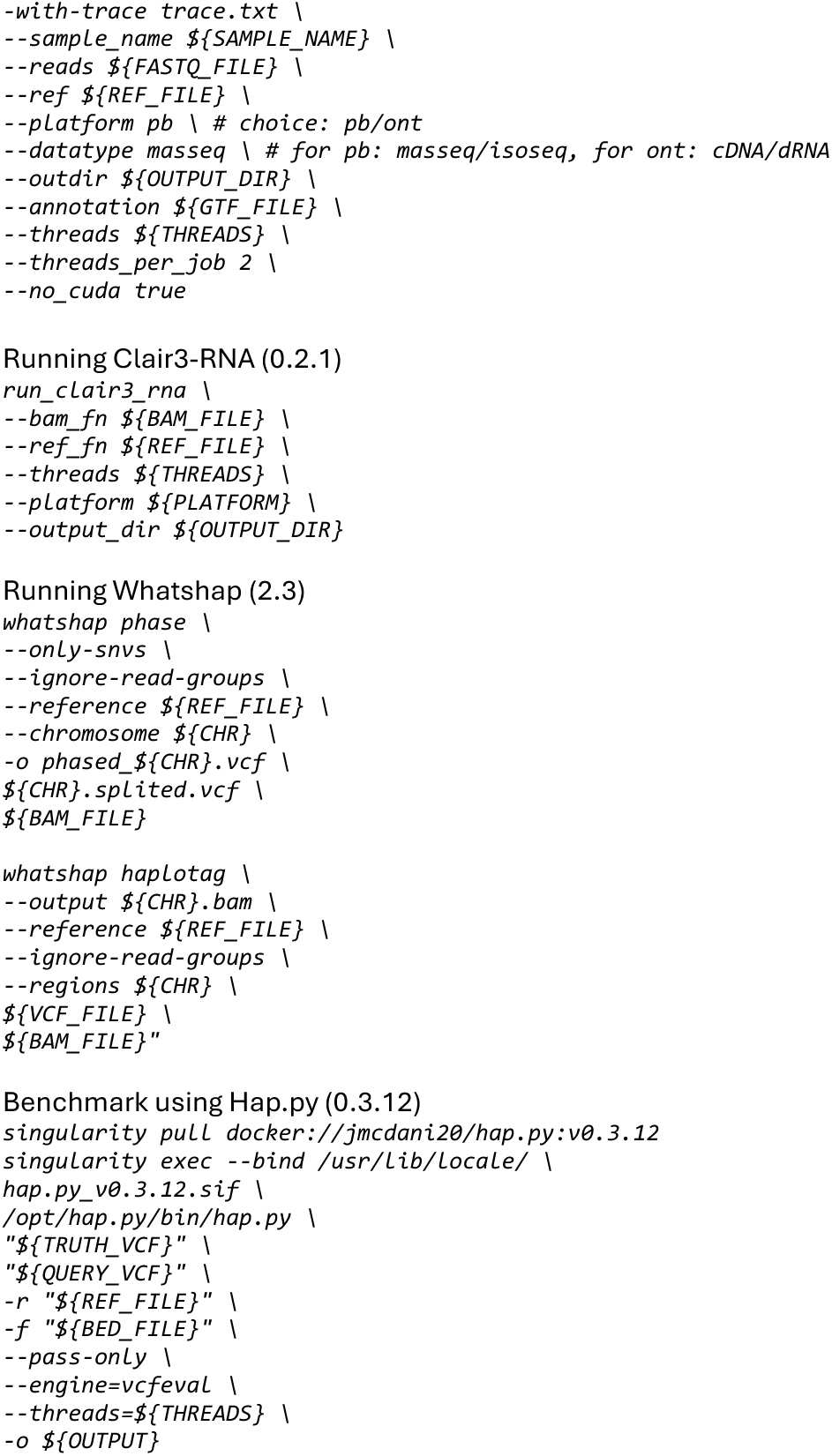

